# Central tendency biases must be accounted for to consistently capture Bayesian cue combination in continuous response data

**DOI:** 10.1101/2021.03.12.434970

**Authors:** Stacey Aston, James Negen, Marko Nardini, Ulrik Beierholm

**Affiliations:** Department of Psychology, Durham University, UK; School of Psychology, Liverpool John Moores University, UK

## Abstract

Observers in perceptual tasks are often reported to combine multiple sensory cues in a weighted average that improves precision – in some studies, approaching statistically-optimal (Bayesian) weighting, but in others departing from optimality, or not benefitting from combined cues at all. To correctly conclude which combination rules observers use, it is crucial to have accurate measures of their sensory precision and cue weighting. Here, we present a new approach for accurately recovering these parameters in perceptual tasks with continuous responses. Continuous responses have many advantages, but are susceptible to a central tendency bias, where responses are biased towards the central stimulus value. We show such biases lead to inaccuracies in estimating both precision gains and cue weightings, two key measures used to assess sensory cue combination. We introduce a method that estimates sensory precision by regressing continuous responses on targets and dividing the variance of the residuals by the squared slope of the regression line, “correcting-out” the error introduced by the central bias and increasing statistical power. We also suggest a complementary analysis that recovers the sensory cue weights. Using both simulations and empirical data, we show that the proposed methods can accurately estimate sensory precision and cue weightings in the presence of central tendency biases. We conclude that central tendency biases should be (and can easily be) accounted for to consistently capture Bayesian cue combination in continuous response data.

## Introduction

One central goal in the study of perceptual decision-making is to understand how an observer makes decisions in the presence of several streams of sensory information. This is a challenge since these streams are always noisy, sometimes biased, and frequently conflicting. To optimise perceptual decision-making, an observer should integrate across all available streams of information that are relevant to the current task. For example, sensory cues from both the visual and auditory domains could be combined to estimate the location of a visually obscured, sound-emitting object. When the observer has prior knowledge about the most likely location of the object, this too should be taken into account. If each piece of information is optimally weighted according to its reliability (or precision), then combination will lead to more precise estimates than if only a single piece of information were used.

There are reports of optimal or near-optimal combination of information for perceptual decision-making in adults both across the senses, with observers optimally integrating visual and haptic cues to depth (Ernst and Banks, 2002) and near-optimally integrating auditory and visual cues to location (Alais and Burr, 2004), and within the senses, with observers optimally integrating disparity and texture gradient cues to slant (Hillis et al., 2002; Knill and Saunders, 2003). This is also true when combining sensory cues and prior knowledge (e.g. Körding and Wolpert, 2004). However, there are also numerous reports of behaviour diverging from optimal (Rahnev and Denison, 2018). To address this, research should explore models of information integration that predict when behaviour is optimal and when it is not (e.g. Laquitaine and Gardner, 2017; Norton et al., 2019).

To do so, it is necessary to create studies where a observer’s sensory precision and/or weighting of the different streams of information can be quantified and compared across conditions. If a researcher is satisfied with using a two-alternative forced-choice method, then techniques for recovering estimates of sensory precision and/or weightings from the data are already well-established in the literature (e.g., Ernst and Banks, 2002). However, if the researcher wishes to track sensory precision and weightings over the course of an experiment, it becomes preferable to allow observers to freely adjust their responses on an appropriately chosen continuous scale. Continuous responses of this type also lead to a more engaging task for the observer, a richer data set that allows for rigorous model comparison to uncover the source of suboptimalities in behaviour, and increased statistical power. This becomes an issue because the analysis techniques available are relatively immature. This paper presents insights and techniques to help improve the analysis of continuous response data.

Specifically, we consider the case of a study with continuous responses and create a reliable method of recovering sensory precision and/or weightings from continuous estimates corrupted by a well-documented bias in perceptual judgements. The bias that we consider is a central tendency bias, where sensory estimates are biased towards the mean of recently seen stimuli. The term goes back to Hollingworth (1910) who demonstrated a central tendency bias for judgements of size. Hollingworth presented observers with a square piece of card, and, after a delay, asked them to choose a card of matching size from a set of standards. The chosen standard for each reference card varied with the series of references shown in any particular block of trials. If the reference was larger than the average reference in the block it was matched to a smaller standard than its true value and vice-versa. Since Hollingworth, central tendency biases have been shown for a range of stimulus types, such as judgements of line length (Ashourian and Loewenstein, 2011; Duffy et al., 2010; Huttenlocher et al., 2000), sweetness (Riskey et al., 1979), facial expressions (Roberson et al., 2007; Corbin et al., 2017), absolute size (Huttenlocher et al., 2000), shades of grey (Huttenlocher et al., 2000), time-interval estimation (Jamieson, 1977; Jazayeri and Shadlen, 2010; Ryan, 2011), and colour (Olkkonen et al., 2014; Olkkonen and Allred, 2014). Clearly, central tendency biases have the potential to corrupt a variety of perceptual judgements.

As we will see, accounting for this type of bias in continuous responses is important for maximising statistical power; it is also important so that the estimate of precision (or variability) that is recovered from the data is truly an estimate of *sensory* precision. In what follows, we first derive how a central tendency bias affects estimates of precision and, hence, how a central tendency bias can reduce the measured gain in precision (what we will call the combination effect) that results from combining information. We then develop an analysis method that accounts for central tendency biases by correcting estimates of precision according to the inferred strength of the bias, thus recovering the combination effect. Through simulation and an application to empirical data, we illustrate how the new method offers a researcher increased statistical power when analysing behavioural data where a central tendency bias is present. For completeness, we also consider the effect of a central tendency bias on inferred cue weightings in continuous responses to conflicting cues, suggesting a method for analysing these types of data that recovers the unbiased weights and, again, we demonstrate the validity of the method through simulations and an application.

## Central tendency biases mask cue combination effects in continuous response data

Formally, given multiple pieces of sensory information, or sensory cues, to a stimulus property, Bayesian cue combination is the optimal way to combine the cues if the goal is to maximise precision. If each cue is represented by a sensory estimate *c*_1_, …, *c_n_* of the true stimulus property, *s*, with Gaussian error distribution, such that 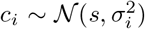, for *i* = 1, …, *n*, the lowest-variance combined estimate is given by 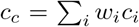, where 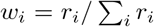, and 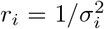 is the reliability of the estimate from the *i^th^* cue. In other words, the optimal way to combine the estimates is to take a reliability-weighted average, placing the most weight on the most reliable piece of information. In the case of two cues, the variance of the combined estimate is 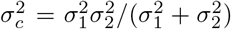, which is lower than the variance of either single cue alone^1^ and results in increased precision when cues are combined.

To capture combination of two cues in a behavioural task, precision is measured using each cue alone and compared with a measurement of precision using both cues together (e.g., Ernst and Banks, 2002). A significant increase in precision using both cues compared to the best single cue illustrates a combination effect and suggests Bayes-like cue combination. If *c*_1_ is the best single cue, the maximum size of the combination effect (or reduction in variance, analogous to the gain in precision) is

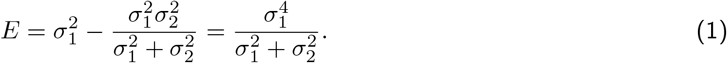

The optimality of the combination can be quantified by comparing measured precision using both cues to the optimal prediction according to precision measured using each cue alone - a test of optimality.

To capture cue combination in continuous responses, a researcher must be sure that they can can recover a cue combination effect. Statistical power for recovering this effect depends on the maximum possible gain in precision (Scarfe, 2020) and the effect may be masked, reducing statistical power, by other aspects of perception and decision-making. For example, the maximum gain in precision (or size of the combination effect) offered by a reliability-weighted average is only guaranteed if the single cue estimates are unbiased (no constant error), have independent Gaussian noise distributions, and are unaffected by other types of perceptual, decision, or response biases. Other authors have addressed cases where single cue estimates are biased (Scarfe and Hibbard, 2011) or where the estimates from different cues have correlated noise distributions (Oruç et al., 2003). In the former case, it is possible to define when optimal cue combination continues to benefit the observer based on the amount of bias between the single cue estimates (Scarfe and Hibbard, 2011). In the latter, it is possible to refine the weights placed on each single cue estimate to account for correlated noise (Oruç et al., 2003). However, to our knowledge, there has been no previous account of the effect that central tendency biases (Hollingworth, 1910; Huttenlocher et al., 2000) can have on combined cue estimates, or how to account for them when measuring a combination effect or computing optimal predictions.

In continuous response data, the response on trial *t* can be modelled as *r_t_* = *c_t_* + *ϵ_t_*, where *c_t_* is the internal estimate on trial *t* (either from a single cue, *c*_1_ or *c*_2_, or both cues, *c_c_*, depending on cue availability), and 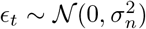 captures any additional response noise (motor noise, for example). The most obvious way to calculate an estimate of variability (or precision) is to take the variance over all errors for a given trial type. For example, if there were *N* trials where only cue 1 was present, the variability of responses using cue 1 could be defined as

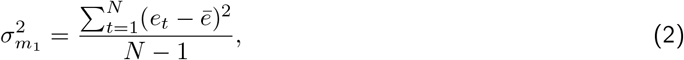

where *e_t_* is the error on trial *t* and we use the subscript *m*_1_ to denote measured variability^2^. Our estimate, 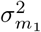, will not be an estimate of 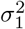, but will be an estimate of 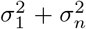, as it encompasses the additional response noise that is added to the internal estimate. However, it can be shown that this preserves the size of the combination effect as^3^

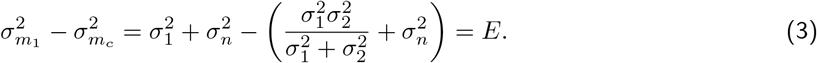

However, if the assumptions made above do not hold (for example, if a central tendency is present) the above equations may not be true, and the size of the combination effect that can be measured from calculating estimates of variability in the way described above may change.

Before we formally derive the effect of a central tendency bias on the size of the combination effect, we consider some hypothetical data to illustrate the effect visually. Suppose six different observers generate the data in Figure 1. Each panel represents data from a separate, hypothetical observer completing a task where they must find a hidden object (along a horizontal axis) using two cues, such an a auditory and visual cue. In the figure, we plot targets on the x axis (the true location of the hidden object) and the observers responses on the y axis. Along the top row, a simple fact is made plain by inspection of the scatterplots: the leftmost observer issues the most precise responses, followed by the middle, followed by the rightmost. The leftmost observer has their responses tightly clustered around the correct target (i.e. a line from the bottom left to the top right of the panel), while the relation between target and response is much looser for the rightmost observer. It is fairly easy to see how, using Equation (2), we could create a measure of variability that would reflect the differences across the top row. This would result in variances of 4, 16, and 36 for the leftmost, middle, and rightmost top figures, respectively. This captures the intuition that was created by simply inspecting the graphs.

**Figure 1:**
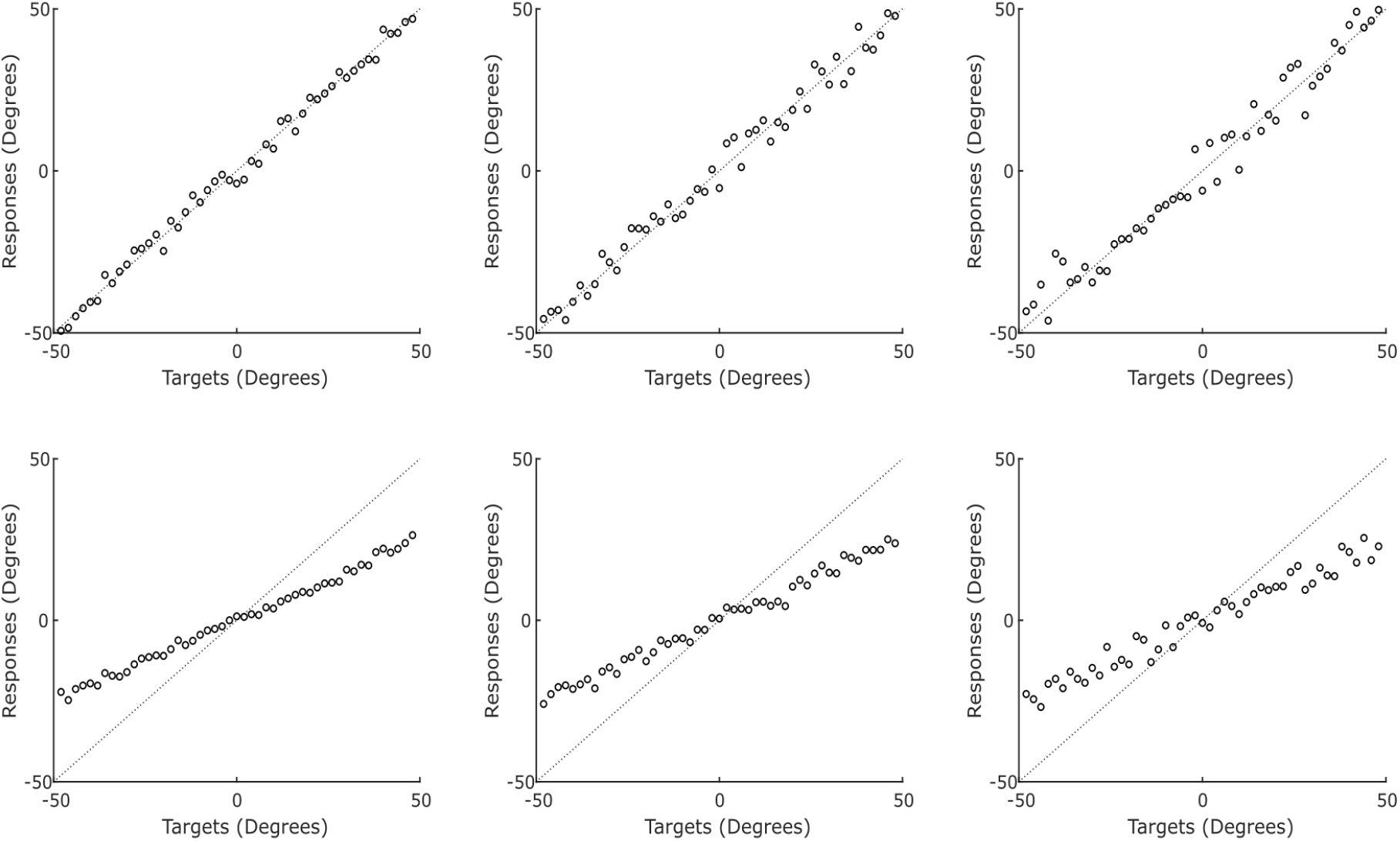
Example responses from six hypothetical observers making perceptual judgements to illustrate the potential issues around calculating sensory variability from continuous responses. In the top row are three observers who do not apply a central tendency bias to their responses, but have varying levels of sensory uncertainty (variabilities of 4, 16 and 36 degrees from left to right). In the bottom row, the observers have the same level of sensory uncertainty as the hypothetical observers above them, but also apply a central tendency bias to their responses.

A problem with over-estimation presents itself when we consider the observers along the bottom row. These observers are subject to a central tendency bias. This is an extreme example for illustration purposes, where the observers display a strong central tendency bias. The observers in these examples equally-weight the sensory information and the central stimulus value. In practice, this moves their estimates 50% of the way towards the central value. We model this as a two-step process, where observers first combine all of the relevant sensory information, forming a *sensory estimate* of the stimulus they must judge, before taking a weighted average of that sensory estimate and the central value to produce their response, or *behavioural estimate*. Our two-step model is supported by empirical findings that central tendency biases arise during reconstruction (or decoding), not encoding of stimulus features (Crawford et al., 2000) and, more importantly, by a model comparison suggesting that multisensory integration precedes a central tendency bias (Murai and Yotsumoto, 2018).

Returning to our hypothetical data, if we again take the variance of the errors, but this time for the observers in the bottom row, it does not capture sensory variability in the desired way. We get a result over 150 for all three observers. In this case, despite the bottom leftmost observer giving very systematic and predictable responses (always near half of the real distance to the center), the raw errors are still very large. For a target at −40, they responded at −19.5 (error of +20.5). For a target at +40, they responded at +21 (error of −19). These systematic biases would become a large part of the calculations. If the variance of the raw errors were used as a measure of sensory variability, it would be an enormous over-estimate that mainly reflected the size of the central tendency bias.

The opposite problem presents itself if we are overzealous in discounting the bias. Instead of looking at the variance of the responses around the correct target, one might look at the variance of the responses around the best fit regression line. However, in the presence of a central tendency bias this will actually give an underestimate of sensory variability, as biasing responses towards the central stimulus value increases the precision of those responses about the biased estimate (likely the reason why the perceptual system adopts a central tendency bias; Huttenlocher et al., 2000). The top leftmost and bottom leftmost observers are generated with the same underlying sensory variability, but after the bottom leftmost observer biases their estimates towards the center, the responses become compressed closer to each other. Subsequently, the variance of the regression residuals is 4 for the top leftmost observer, but only 1 for the bottom leftmost observer. The remaining pairs have 1/4 of the variance in the right column as well.

We propose a solution that balances between the over-estimate given by the variance of the errors and the under-estimate given by the variance of the regression residuals. In short, our solution regresses responses, *r_t_*, on targets, *s_t_*, to estimate *α* and *β* such that *r_t_* = *βs_t_* + *α* + *ϵ_t_*. We propose that the variance of the residuals, *ϵ_t_*, divided by the squared slope of the regression line, *β*^2^, is the best estimate of sensory variability that can be recovered from the data. In our example above, this assigns a sensory variance of 4, 16, and 36 to both members of the leftmost, middle, and rightmost columns, respectively. As desired, this is neutral to the fact that members of the bottom row have a central tendency bias and members of the top row do not, capturing the underlying sensory variance that was used to generate the examples. Below, we justify this in detail with a formal mathematical treatment of the issue.

Following our two-step model, responses corrupted by a central tendency bias can be modelled as a weighted average of (combined) sensory estimates, *c_i_*, and the center (or mean) of the stimulus distribution, *μ*. Single and combined cue behavioural estimates are then

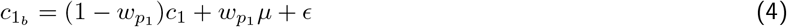

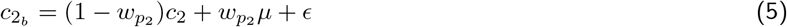

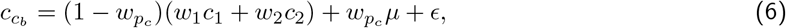

where *w_p_i__* is the weight placed on the center of the stimulus distribution relative to the sensory estimate. The variabilities of the behavioural estimates, *c*_1_*b*__, *c*_2_*b*__, and *c_c_b__*, are 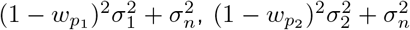, and 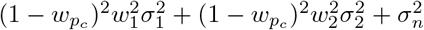, respectively. This reduces the measurable size of the combination effect as

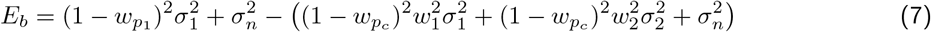

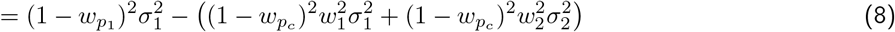

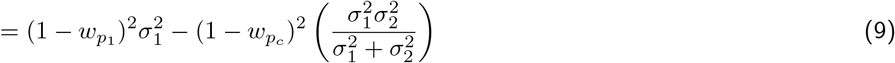

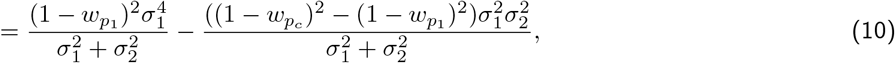

and, *E_b_* < *E* as, in the case of a non-negligible weight on the center of the stimulus distribution (i.e. *w*_*p*_1__, *w_p_c__* > 0), (1 – *w*_*p*_1__)^2^ < 1 and *w*_*p*_1__ ≥ *w_p_c__* implies (1 – *w*_*p*_1__)^2^ ≤ (1 – *w_p_c__*)^2^, so the first term is smaller than *E* and the second term is greater than or equal to zero. Empirical evidence (Olkkonen et al., 2014; Olkkonen and Allred, 2014) and Bayesian models of central tendency biases (Huttenlocher et al., 2000; Jazayeri and Shadlen, 2010; Cicchini et al., 2012; Sciutti et al., 2014; Krjgel et al., 2020) imply that the strength of the bias (or weight on the central value) will increase with increasing sensory uncertainty or variability. This satisfies our above assumption that *w*_*p*_1__ ≥ *w_p_c__*, as sensory uncertainty should be reduced following reliability-weighted averaging of sensory information.

We have already alluded to a distinction between *sensory variability* (or *sensory precision*) and *behavioural variability* (or *behavioural precision*), but we will make that distinction explicit here. As central tendency biases appear to be introduced after the sensory estimate is formed (Crawford et al., 2000; Murai and Yotsumoto, 2018), comparing the variability of behavioural responses across different experimental conditions (e.g. when observers use their best single cue compared to multiple cues) will not reflect the reduction in variance, or gain in precision, that is afforded from taking a reliability-weighted average of sensory information, or the underlying combination effect *E* (Equation 1). Instead, an analysis that uses behavioural variabilities may fail to find a combination effect because the behavioural gain in precision, *E_b_*, will be smaller than the sensory gain, *E*, if estimates are centrally biased (Equations 8–10)^4^. We would like a method for analysing the behavioural responses that recovers a measure of sensory rather than behavioural precision; that is what we offer here.

## Accounting for central tendency biases in continuous responses to recover cue combination effects

Previously, we modelled continuous responses as *r_t_* = *c_t_* + *ϵ_t_*, where 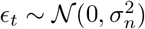. When a central tendency bias is present, *c_t_* is replaced with *c_t_b__* and the response on trial *t* becomes *r_t_b__* = *c_t_b__* + *ϵ_n_*. Assuming that 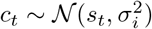, we can rewrite this as

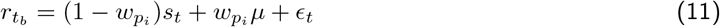

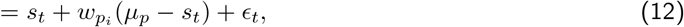

where *i* = 1, 2, or *c* depending on trial type. For each pairing of trial type and stimulus, *s_t_*, the biasing part of the equation, *w_p_i__*(*μ* – *s_t_*), is a fixed constant error. This constant error must be calculated for each stimulus independently and subtracted from all responses to that stimulus before Equation (2) can be applied to calculate behavioural precision. Alternatively, the experimenter could calculate an estimate of behavioural precision by regressing responses on the true stimulus values and taking the variance of the fitted residuals. We will adopt this approach here as it can always be used while the former is only possible if each stimulus value is tested multiple times.

To calculate an estimate of sensory precision, we must consider Equation (11). The second term in this equation is fixed for each trial type, allowing us to model all responses for a given trial type as *r_t_b__* = *βs_t_*+*α*+*ϵ_t_*, making it clear that the constant term, *w_p_i__μ*, only changes the intercept of the regression line relating the responses to the true stimulus values. Importantly, the fitted coefficient, *β*, which represents the slope of the line and captures the level of bias (lower values for *β* imply less weight on the stimulus and, thus, more bias), is an estimate of (1 – *w_p_i__*). Since our measure of behavioural precision is an estimate of 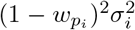, we can divide by the square of the fitted coefficient, *β*^2^, to gain an estimate of the underlying sensory precision.

Note that if *β* is approximately 1, as is the case when the strength of the bias is negligible or there is no bias at all (*w_p_i__* = 0), sensory precision is identical to behavioural precision. This also makes it clear that if *β* is greater than one its value should be ignored. In practice, when applying the analysis method to the simulated and empirical data later in the text, we account for this by only applying the correction if the fitted coefficient, *β*, is significantly less than one at the 5% significance level. This makes the proposed analysis method flexible, as a central bias in the data will only be accounted for if it is present.

We note that the measure of behavioural precision (the estimate before the correction) is an estimate of 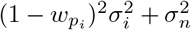, and so our estimate of sensory precision is an estimate of 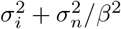. This changes the maximum size of the combination effect we can detect as

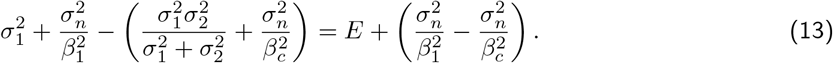

Under the assumption that the central tendency bias increases with decreasing sensory precision (Olkkonen et al., 2014; Olkkonen and Allred, 2014; Huttenlocher et al., 2000; Jazayeri and Shadlen, 2010; Cicchini et al., 2012; Sciutti et al., 2014; Krügel et al., 2020), *β_c_* > *β*_1_, and the second term on the right side of Equation (13) is greater than zero, increasing with increasing *σ_n_*. In other words, there are certain circumstances where the additional noise, *σ_n_*, and the difference between the strength of the bias when using the best single cue alone, *β*_1_1__, and both cues together, *β*_1_*c*__, are large enough that this method not only recovers statistical power for detecting a combination effect, but can enhance the effect and increase statistical power over and above what recovery of raw sensory precision would allow.

We argue that as the intention of the proposed analysis is only to uncover potentially hidden combination effects in continuous response data, not to provide an exact estimate of the size of the combination effect, then this is a positive. The reader may of course worry that it will lead to reports of combination effects that are not actually there. This is not the case if the strength of the central tendency bias is determined by the level of sensory precision. Without an improvement in sensory precision using multiple cues compared to the best single cue (i.e. *E* = 0), *β_c_* = *β*_1_ and the second term on the right side of Equation (13) is exactly zero.

## An illustration with simulated continuous responses

We simulated data from a optimal Bayesian model of cue combination, where an observer takes a reliability-weighted average of multiple cues to maximise their gain in precision and, hence, their combination effect. However, the observers that we simulated were subject to a central tendency bias (Equations (4)–(6)). For these simulations, we assumed that all stimulus values were in the range 0-1 (other ranges can easily be mapped here) and used targets, *s*, of 0.15 to 0.85 in steps of 0.02. We generated five responses for each target and each trial type (cue 1 only, cue 2 only, both cues) for every observer that we simulated.

The simulated observers differed in their cue reliability ratios and the strength of their central tendency bias. We fixed the reliability of the best cue, cue 1, so that *σ*_1_ = 0.01, and varied the reliability ratio so that *σ*_2_/*σ*_1_ was one of twenty log-spaced values between 1 and 10. To vary the strength of the central tendency bias, we varied the weight that each observer placed on the center of the stimulus range in Equation (6) by setting *w_p_c__* equal to one of twenty log-spaced values between 0.01 and 1. As the strength of the central tendency bias can vary within an observer according to the reliability of the sensory evidence, we defined *w*_*p*_1__ and *w*_*p*_2__ as 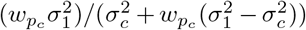 and 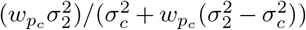, respectively, so that the weight on the center of the stimulus range increased as the reliability of the sensory information decreased. These equations are chosen so that our simulations are in line with Bayesian accounts of central tendency biases (see Discussion). For each pairing of reliability ratio and strength of the bias, we simulated 1000 data sets at five different additional noise levels, *σ_n_* = 0,0.005,0.01, 0.015, or 0.02, i.e. no additional noise, or additional noise at 0.05%, 0.1%, 0.15%, or 0.2% of the stimulus range.

We calculated statistical power for detecting a gain in precision, or a combination effect, when multiple cues were present by comparing response variability with both cues to response variability with the best (lowest variability) cue alone using a one-tailed Wilcoxon-Signed Rank Test (*α* = 0.05) for three different sample sizes (*n* = 10, 20, or 30). To do so, for each pairing of reliability ratio and bias strength, each additional noise level, and each sample size, we took 100 randomly sampled data sets of size *n* from the 1000 relevant simulations. For each randomly sampled data set, we calculated estimates of variability when using each single cue alone and when using both cues for each simulation in that set. To calculate the estimates we fit a regression line to the responses with true stimulus value as the independent variable. The variance of the fitted residuals provides an estimate of *behavioural precision*, that does not account for any central tendency bias in the responses. Alternatively, dividing the variance of the fitted residuals by the square of the fitted slope ”corrects-out” any precision that is gained by applying a central tendency bias to recover an estimate of *sensory precision*, the value that is of interest to us.

Figure 2 shows power for detecting a combination effect for different pairs of reliability ratio and bias strength, at different additional noise levels, and with different sample sizes using the estimates of behavioural precision. It is clear from this figure that increasing the reliability ratio, the strength of the bias, or the level of additional noise decreases statistical power for detecting a combination effect. Increasing the sample size recovers some of the power that is lost by the increase in reliability ratio or additional noise, but does not help to recover power that is lost by the increase in the strength of the central tendency bias. Indeed, once the bias surpasses 0.3, or more than a 30% shift towards the central stimulus value, our ability to detect a combination effect in these simulations is effectively lost, regardless of reliability ratio, level of additional noise, or sample size. However, Figure 3, which shows power for detecting a combination effect when using our proposed measure of sensory variability that accounts for a central tendency bias, tells a different story. In this figure, power is still lost by increasing the reliability ratio or the level of additional noise, with both partially counteracted by increasing the sample size, but power does not decrease as the strength of the bias increases. This is because our proposed analysis method, that calculates the measure we refer to as sensory variability, accounts for the central tendency bias to recover the sensory precisions. Figure 4 directly compares statistical power using the two measures (behavioural and sensory precision) by subtracting the values depicted in Figure 2 from those depicted in Figure 3. This figure shows that using our proposed analysis method to account for a central tendency bias increases statistical power regardless of additional noise level or sample size when the strength of the bias is greater than 0.3 and the ratio of the reliabilites is less than 5. As the level of additional noise increases, our proposed method begins to increase power for lower bias strengths.

**Figure 2:**
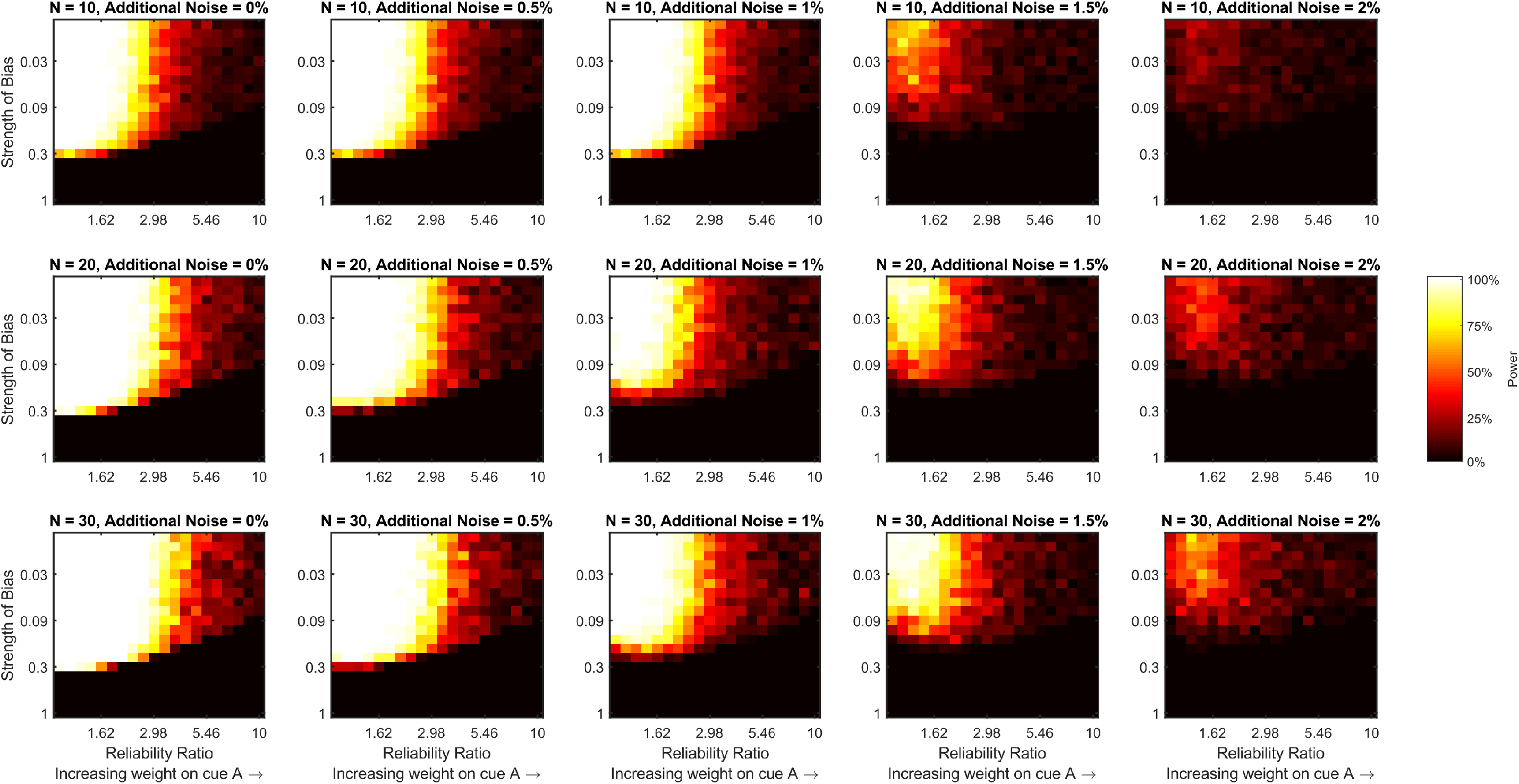
Statistical power for detecting a combination effect using measures of variability (or precision) that do not account for a central tendency bias. In the text, we refer to this as a measure of behavioural variability, calculated by taking the variance the fitted residuals, *ϵ_t_*, when regressing responses, *r_t_*, on targets, *s_t_*, to estimate *α* and *β* such that *r_t_* = *βs_t_* + *α* + *ϵ_t_*. Each panel of the figure is a heat map that shows how statistical power (estimated by bootstrapping the simulated data sets) varies with the ratio of the two cues’ reliabilities and the strength of the central bias for a fixed level of additional noise and a fixed sample size. The level of additional noise varies across the columns of the figure and sample size varies across rows.

**Figure 3:**
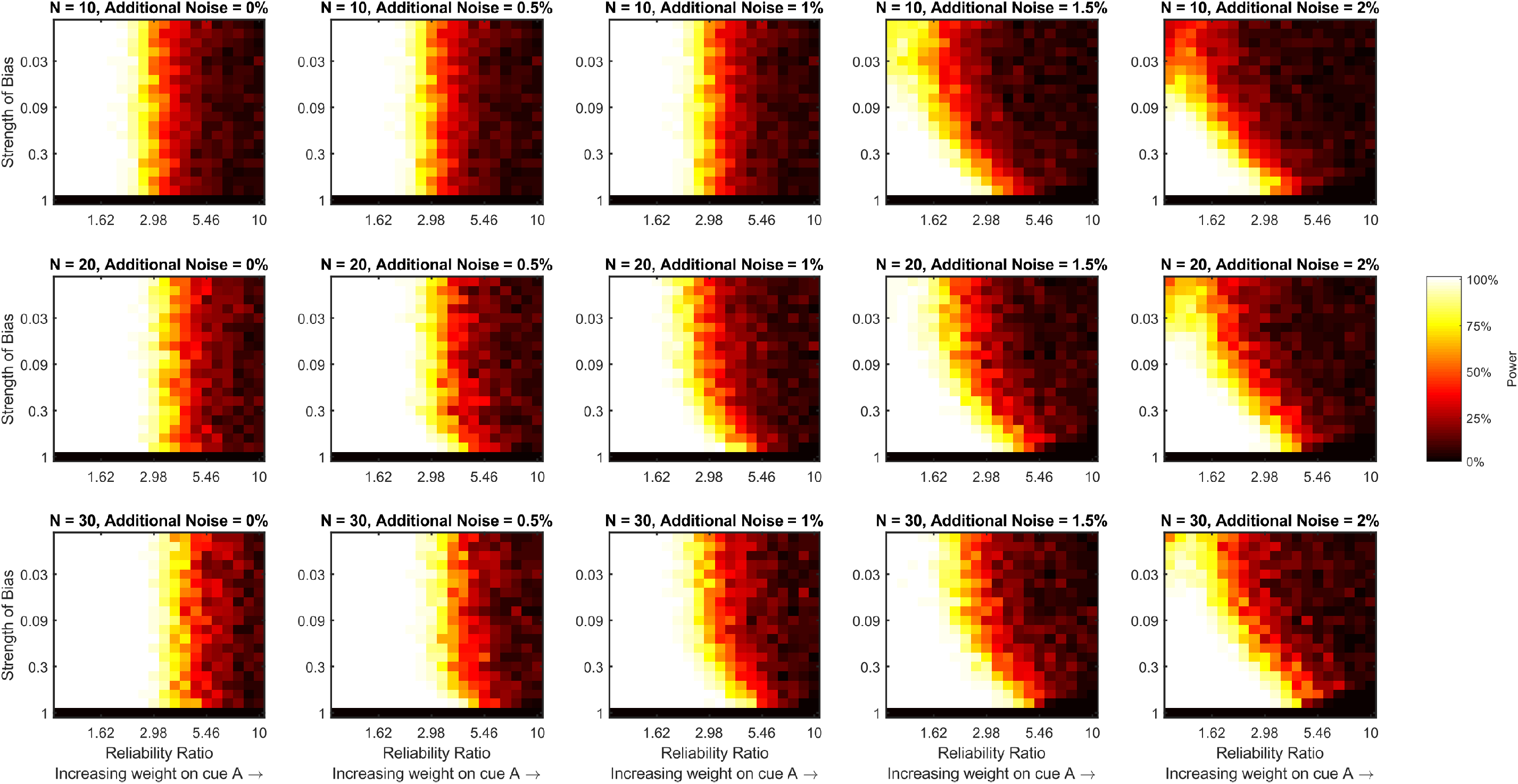
Statistical power for detecting a combination effect using the measure of variability (or precision) introduced here that accounts for a central tendency bias. In the text, we refer to this as a measure of sensory variability, calculated by dividing the variance the fitted residuals, *ϵ_t_*, by the square of the fitted gradient, *β*^2^, when regressing responses, *r_t_*, on targets, *s_t_*, to estimate *α* and *β* such that *r_t_* = *βs_t_* + *α* + *ϵ_t_*. Each panel of the figure is a heat map that shows how statistical power (estimated by bootstrapping the simulated data sets) varies with the ratio of the two cues’ reliabilities and the strength of the central bias for a fixed level of additional noise and a fixed sample size. The level of additional noise varies across the columns of the figure and sample size varies across rows.

**Figure 4:**
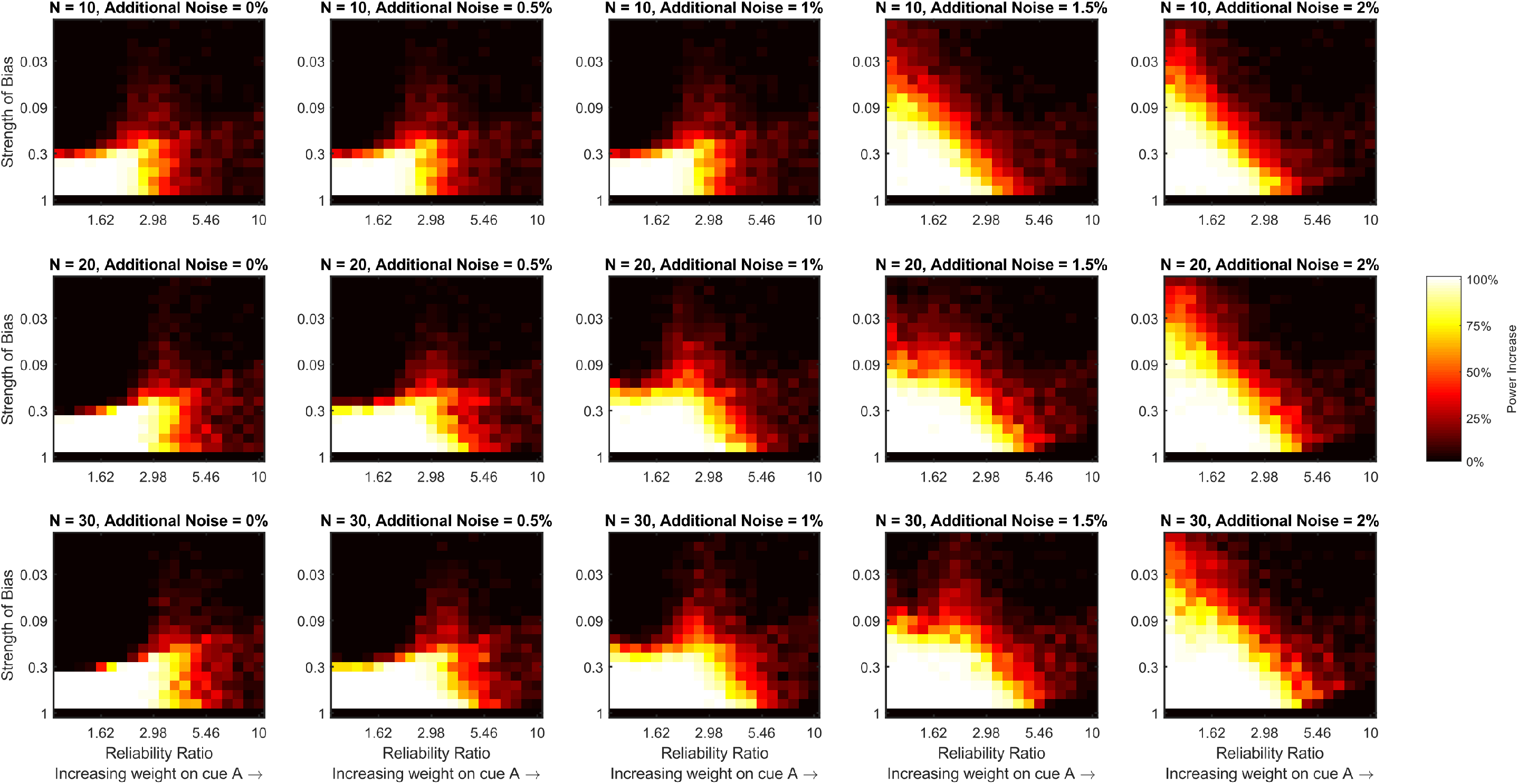
The increase in statistical power for detecting a combination effect gained by using the measure of sensory variability introduced here (that accounts for a central tendency bias) rather than, what we refer to as, a measure of behavioural variability (that does not account for a central tendency bias). Each panel of the figure is a heat map that shows how the increase in statistical power (estimated by subtracting the values depicted in Figure 2 from those depicted in Figure 3) varies with the ratio of the two cues’ reliabilities and the strength of the central bias for a fixed level of additional noise and a fixed sample size. The level of additional noise varies across the columns of the figure and sample size varies across rows.

## An example with empirical two-cue continuous response data

Here, we demonstrate the effect of a central tendency bias on estimates of the combination effect in previously published data. We re-analysed the data from Experiment 1 in an article recently published in *Cognition* (Negen et al., 2019). This study presented 77 7-10 year old children with audio, visual, or audio-visual cues to a horizontal location. Observers pointed and clicked on the perceived source of the stimulus. We adopt the same observer exclusion criteria as in the original publication, leaving us with 68 observers for our analysis. We will calculate three separate estimates of the combination effect from the data. The first will be calculated by estimating precision using each single cue alone and both cues from the variance of the *raw* errors (response-target), as in the original paper. We will refer to this as a measure of *raw precision*. This method does not account for the constant error that varies across location in the presence of a central tendency bias, or the extra precision that is gained from applying a central bias over and above that offered by combining the sensory information. The second estimate will be calculated by taking the variance of the residuals about a fitted regression line that regresses responses on targets; defined as a measure of *behavioural precision* earlier in the text. This measure accounts for the varying constant error introduced across location by a central tendency bias but does not account for the added precision. The third estimate, our measure of *sensory precision*, will be calculated by dividing the variance of the fitted residuals by the square of the target’s coefficient from the fitted regression line. This estimate “corrects-out” both the constant error and the added precision from the central bias.

Before we look at how the three measures of precision differ, we can consider the average slope of the regression lines to see if there was evidence of a significant central tendency in these data. If the average slope is significantly less than 1, then the average weight on the central value of the stimulus range is significantly greater than zero, and there is a significant central tendency bias. The average weight on the cue(s) was significantly less than one in the visual trials (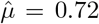, *t*(67) = −14.59, *p* < .001), the audio trials (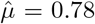, *t*(67) = −5.81, *p* < .001), and the audio-visual trials (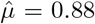, *t*(67) = −6.44, *p* < .001), illustrating a significant central tendency bias for all trials types. Moreover, the average weight on the cue differed significantly across trial types (*F*(2,134) = 11.96, *p* < .001), with most weight placed on the cue in audio-visual trials and least in the visual trials. This is in line with Bayesian theories of a central tendency bias, where the centre of the stimulus range receives more weight as cue uncertainty increases (Olkkonen et al., 2014; Olkkonen and Allred, 2014; Huttenlocher et al., 2000; Jazayeri and Shadlen, 2010; Cicchini et al., 2012; Sciutti et al., 2014; Krügel et al., 2020).

Given the presence of a central tendency bias in the data, we found varying sizes of the combination effect using the three different methods to estimate precision. A one-tailed sign-rank test comparing each observer’s precision using both cues to that using their best single cue alone found a significant combination effect regardless of the calculation that was used to estimate variance (raw: *z* = 5.15, *p* < .001; behavioural *z* = 2.04, *p* = .02; or sensory *z* = 4.55, *p* < .001). However, the size of the effect varies widely depending on the analysis that is used. Using the raw variances, Cohen’s effect size was *d_r_* = 0.72, with 78% of observers showing a gain in precision using both cues. Using behavioural variance, Cohen’s effect size was *d_b_* = 0.27, with only 60% showing the effect. Using the sensory variance, Cohen’s effect size was *d_s_* = 0.41, with 82% showing the effect. This clearly illustrates the point we have already made through an informal example, formal derivation, and simulation: a central tendency bias reduces statistical power to detect a combination effect if only the varying constant error across location that is introduced by it is accounted for (in the behavioural variance case), or could cause the effect to be over-estimated if only the raw errors are considered. Using the sensory variance lands the estimate of effect size somewhere in the middle of those two estimates, recovering the underlying estimates of sensory precision if the process model follows that which we have laid out above.

## Central tendency biases lead to underestimates of cue weightings in continuous responses to conflicting cues

Thus far, we have only considered how a central tendency bias may alter estimates of precision in a scenario where a Bayes-optimal observer is combining two unbiased cues that are not in conflict. However, in experiments that study cue combination, the researcher is often interested in measuring how two cues are weighted relative to each other. To do this, a researcher designs an experiment so that the two cues indicate conflicting target values (e.g., Ernst and Banks, 2002). If the conflict is kept small enough, the Bayes-optimal observer should continue to take the reliability-weighted average of the two cues to maximise perceptual precision. In the absence of a central tendency bias, either (a) a multiple regression of responses on predictors cue 1 and cue 2, or (b) a simple regression of response bias relative to cue 2 (response-cue 2) on conflict in the direction of cue 1 (cue 1-cue 2), gives an estimate of the weight placed on each cue. Using (a) provides a direct estimate of both weights (weight on cue 1 and on cue 2). Using (b) only directly estimates the weight on cue 1, but contains the implicit information that the two weights sum to unity. This is not the case in (a) unless constraints are used when performing the multiple regression. Regardless, when a central tendency bias is present, the fitted weight on cue 1 is an estimate of (1 − *w*_*p*_1__)*w*_1_ rather than *w*_1_, leading to an underestimate of the weight on the cue. To avoid this issue, one must consider a model of responses that allows for a central tendency bias (Equation (6)) and estimate the parameters accordingly. Here, we show how this approach leads to better estimates of the true sensory cue weightings.

To do so, we again simulated data from a Bayes-optimal model of cue combination that is subject to a central tendency bias (see Equations (4)–(6)). Again, we assumed that all stimulus values were in the range 0-1. However, in this set of simulations, the target indicated by the two cues differed, or was in conflict. For illustration purposes, the Bayes-optimal observers that we simulate combine the cues regardless of the level of conflict between them. We created conflicting cue pairs by taking all possible pairings of target values 0.4 to 0.6 in steps of 0.025 and allowing all reversals of these pairings. We generated ten responses for each pair for every observer that we simulated.

As before, the simulated observers differed in their cue reliability ratios and the strength of their central tendency bias with the reliability of the best cue fixed at *σ*_1_ = 0.05. As the procedure we used for fitting the central tendency bias model was computationally intensive, we chose to simulate fewer reliability ratios, only simulating *σ*_2_/*σ*_1_ as one of five log-spaced values between 1 and 4.18. Similarly, we varied the strength of the central tendency bias, *w_p_c__*, as before but allowed it to be only one of ten log-spaced values between 0.01 and 0.7. For each pairing of reliability ratio and strength of the bias, we simulated 100 data sets at three different additional noise levels, *σ_n_* = 0.01, 0.015, or 0.02.

For each set of simulated responses, we calculated an estimate of the weight on cue 1 without taking into account a central tendency bias by regressing response bias relative to cue 2 (response-cue 2) on conflict in the direction of cue 1 (cue 1-cue 2). We will refer to this as the behavioural weight. We also calculated an estimate of the weight on cue 1 under the assumption of a central tendency bias. To estimate parameters of the model (Equation (6)) for each simulated data set, we used a Gibbs Sampler (JAGS; Plummer, 2003) implemented in MATLAB using the MATLAB-to-JAGS interface matjags.m. We ran three independent chains, discarding the first 100 samples of each chain as burn-in, and recording 1000 samples after the burn-in period, thinned by recording only every 5^th^ sample. Both fitted weights (*w*_1_ and *w_p_c__*) were initialised at 0.5 in all chains. The standard deviation of the additional noise (*σ_n_*) was initialised at 0.01. The resulting estimates were taken as the mean of the expected values from the three chains. We refer to this estimate of the weight on cue 1 as the sensory weight.

Figure 5 illustrates that the weight on the best cue is underestimated when we estimate weights from the simulated data using a method that does not allow for a central tendency bias (the behavioural weight) if the reliability ratio is large enough and the central bias is strong enough. The size of the underestimation does not depend on the level of additional noise. However, when we estimate weights from the simulated data using a method that allows for a central tendency bias (the sensory weight), the error in the estimated weight is zero (on average) regardless of the reliability ratio or strength of the bias. Thus, by estimating cue weightings according to our proposed methods, the estimated weights will remain good estimates of the true weightings of Bayesian observers, even if the observers apply a strong central bias to their responses.

**Figure 5:**
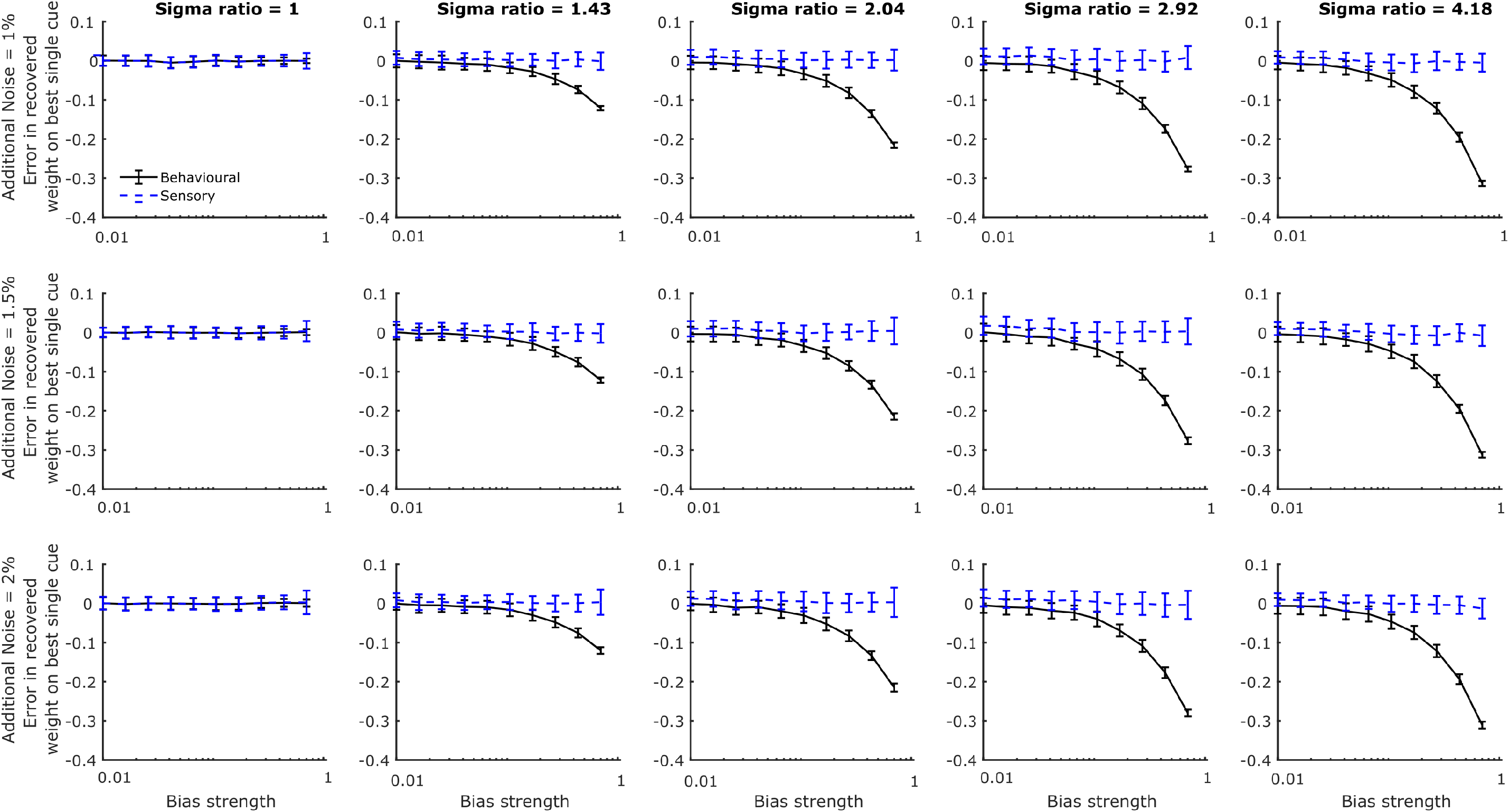
The error in the recovered weight on the best cue estimated from simulated data using a model of responses that accounts for a central tendency bias (the sensory weight) and using a model that does not account for a central tendency bias (the behavioural weight). Each panel of the figure shows how the error in the recovered sensory and behavioural weights varies with the strength of the central bias (on a log scale) for a fixed level of additional noise and a fixed reliability ratio. The reliability ratio varies across the columns of the figure and the level of additional noise varies across rows. Error bars are ±1 standard deviation.

## An example with empirical conflicting two-cue continuous response data

Here, we demonstrate the effect of a central tendency bias on estimates of cue weights in data previously presented at an international conference (Aston et al., 2020). We re-analysed the data from this study where 30 adult observers were presented with intrinsically noisy audio, visual, or conflicting audio-visual cues to horizontal location and freely moved a mouse to respond. We ignored the trials featuring extrinsic noise.

We will first test the hypothesis that observers significantly mis-weight the cues without allowing for a central tendency bias to be present in any of the analysis. To do so, we estimate the optimal weights according to measures of behavioural precision using the audio and visual cues alone. We compare the estimates of the optimal weights to measures of the behavioural weight placed on each cue in the audio-visual trials. These are found by regressing response bias relative to cue 2 (response-cue 2) on conflict in the direction of cue 1 (cue 1-cue 2). We will then test the same hypothesis using weights that are estimated from the data under the assumption of a central tendency bias. Here, the optimal weights are estimated according to measures of sensory precision using each cue alone. They are compared to estimates of the weights used by observers that come from modelling responses as in Equation (6) and estimating the parameters of this model according to the procedure described in the previous section.

In calculating estimates of behavioural and sensory precision, we regress responses on targets, with the slope of the line providing a measure of the strength of any central tendency bias that is present. Similar to the previous empirical data that we considered, we find evidence of a significant central tendency bias (slope less than 1) using both the auditory and visual cues alone (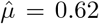, *t*(29) = −7.49, *p* < .001 and 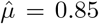, *t*(29) = −24.15, *p* < .001, respectively). Our measure of bias strength for the audio-visual cue condition comes from the model fit (it is the fit parameter *w_p_*) and is also significant (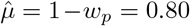, *t*(29) = −10.16, *p* < .001). Taken together, these results point towards the presence of a central tendency bias.

Subsequently, we find a significant difference between the behavioural and sensory weight estimates, *z* = 2.36, *p* = .019. We find no evidence that observers significantly mis-weight the cues regardless of whether we use the behavioural or sensory weights to test the hypothesis. However, the test is closer to significance when behavioural weights are used (*z* = 1.76, *p* = .079, 67% underweight the cues) than when sensory weights are used (*z* = −1.45, *p* = .147, 43% underweight the cues), as expected according to our simulations if these observers are in fact taking a reliability-weighted average of the cues.

## Discussion

Central tendency biases have been demonstrated for a wide range of perceptual judgements about stimuli that come from a predefined range or that are uniformly distributed (Ashourian and Loewenstein, 2011; Duffy et al., 2010; Huttenlocher et al., 2000; Riskey et al., 1979; Roberson et al., 2007; Huttenlocher et al., 2000; Jamieson, 1977; Jazayeri and Shadlen, 2010; Ryan, 2011; Olkkonen et al., 2014; Olkkonen and Allred, 2014). Subsequently, central tendency biases are likely to be found in tasks that aim to capture cue combination. We have explained how this type of bias (a bias towards a specific point in the stimulus-response space) results in increased behavioural precision, which does not capture the measure of sensory precision that would reflect a sensory cue combination effect. Put simply, the analysis method we have proposed ”corrects-out” the added gain in precision that the central tendency bias affords to recover the underlying sensory measure of precision that a cue combination researcher is most likely to be interested in.

Our suggestion for dealing with central tendency biases when calculating the weight that is placed on each cue is even more direct than the correction for recovering sensory precision. To calculate sensory rather than behavioural weights we simply suggest that a researcher allows for a central tendency bias in the response model. If a central tendency bias is not present in the data, then the parameter representing the strength of the bias will be small and the researcher will see no difference in the calculated weights using either a model that does or does not allow for a central tendency bias. However, by adopting this approach the bias will be accounted for if it is there.

### The limiting factor in tests of optimality

By carefully working through the details of a Bayes-optimal observer model completing a continuous cue combination task, we have shown that researchers are limited in their ability to test for an optimal gain in precision when combining cues if the responses are corrupted by additional sources of noise and/or a central tendency bias is present (see footnote 3/4 and Equation (13) showing our method does not recover the exact combination effect in the case of non-negligible additional response noise).

This is an important observation in a research climate that is actively debating the optimality of perception and decision-making processes (Rahnev and Denison, 2018). Clearly, we must be careful in setting the optimal criteria that we compare behaviour against. In footnote 3, we show that regardless of whether a central tendency bias is present or not, non-negligible additional response noise will lead to a definition of optimal precision that is impossible for observers to achieve. We encourage researchers to think carefully about models of behaviour across a variety of tasks where additional sources of noise, or other common features of perception such as a central tendency bias, may lead to optimal predictions that do not reflect what can actually be achieved by the observers.

Quantitative computational models can be used to isolate specific factors in a decision making framework (Odegaard et al., 2019; Jones et al., 2019), including central biases, but this is as of yet not a method that is readily available for experimental researchers.

### The proposed analysis is consistent with Bayesian accounts of a central tendency bias

Huttenlocher and colleagues developed the Category Adjustment Model (CAM) as an account of central tendency biases (Huttenlocher et al., 2000). The CAM proposes that stimulus properties are coded hierarchically at two levels. One is defined as the fine-grained stimulus level (a sensory estimate), that provides an inexact (noisy) but unbiased estimate of the stimulus property. The second is the category level, where the stimuli is encoded as a category prototype, providing a more stable but biased estimate. The combination is a reliability-weighted average, identical to the way estimates from multiple cues are integrated in the Bayes-optimal cue combination model. Later accounts of central tendency biases take a similar approach, expressing central tendency biases within a Bayesian framework (Jazayeri and Shadlen, 2010; Cicchini et al., 2012; Sciutti et al., 2014; Krügel et al., 2020). Recent publications question the ability of CAM, or Bayesian models more generally, to explain central tendency biases in perceptual judgements, suggesting instead that central tendency biases are the manifestation of a recency bias rather than a central bias (Duffy and Smith, 2018, 2020b) but the debate is ongoing (Crawford, 2019; Duffy and Smith, 2020a).

Our approach to modelling central tendency biases is consistent with the Bayesian framework if we express the category level representation as a Bayesian prior that encodes the mean of the stimulus range as the category prototype (the expected value of the prior) and changes in (un)certainty about category membership (the variability of the prior) across the stimulus range. There are two ways we can frame the combination of stimulus and category level information in the Bayesian framework. The options are to combine each individual cue estimate with the category prototype during encoding, before the individual estimates are combined, or to bias the combined estimates during decoding. Crawford et al. (2000) found that estimates of line length in the Müller-Lyer illusion were only subject to a central tendency bias when estimates were made after a delay, interpreting this as evidence that the central tendency bias occurs during decoding to account for uncertainty. This is supported by the findings of Olkkonen and Allred (2014) who found that adding either internal or external noise increased the magnitude of a central tendency bias for estimates of colour, also suggesting that the bias is introduced during reconstruction from memory. Our model is in line with these results where the bias towards the mean of the stimulus range is applied after combination, while being agnostic regarding the mechanism.

### Extending the method to model central tendency biases in tasks where one of the cues is prior knowledge

Closely related to cue combination tasks are tasks where observers are encouraged to combine a single cue with prior knowledge to optimise perception and decision-making. Everything we have discussed could be framed in terms of a cue-prior combination task rather than a cue-cue combination task by letting all parameters that represent one of the cues in each of our equations represent the prior knowledge instead. For example, many of the studies focusing on integration of sensory and prior information require observers to estimate the location of a hidden target, on a continuous scale, using an uncertain sensory cue and prior knowledge of the target distribution (e.g., Berniker et al., 2010; Tassinari et al., 2006; Vilares et al., 2012; Chambers et al., 2018; Kiryakova et al., 2020; Bejjanki et al., 2016). In some cases, when the target distribution is centered on the middle of the stimulus-response range, it would be difficult to parse how much of the central bias is due to a reliance on prior knowledge or a central tendency bias. However, if the center of the target distribution is offset from the center of the stimulus-response range then it may be possible to separate the two factors by using a response model that allows for a central bias as we have done for the two-cue case.

## Conclusion

We have presented a new method of analysis that accounts for central tendency biases in continuous response data to recover estimates of what we define as sensory rather than behavioural precision. We have also suggested a method for modeling a central tendency bias when estimating cue weightings from continuous responses. We illustrated theoretically, through simulations, and through empirical examples, that the application of the new method is necessary and that these biases exists in previously presented cue combination data. The analysis for computing sensory precision is simple and easy to apply. It boils down to dividing the variance of the residual behavioural errors, a measure of behavioral precision, by the squared slope of a regression line that regresses the responses on the true stimulus values. It is also flexible in the sense that a bias in the data will only be accounted for if it is present. If there is no central tendency bias, then the fitted slope value will be approximately 1, and sensory precision is approximately equal to behavioural precision. Thus, the analysis can always be used, even if the data is free of a central tendency bias.

## Acknowledgements

This project has received funding from the European Research Council (ERC) under the European Union’s Horizon 2020 research and innovation programme (grant agreement No. 820185) and a Leverhulme Trust Research Project Grant (RPG-2017-097).

## Open Practices Statement

The MATLAB code to run all simulations and the the two sets of empirical data that we analyse are available here: osf.io/zxmpf.

## Footnotes

1 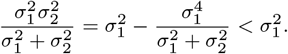

2 Note that we make an assumption throughout that the text that single cue estimates are unbiased in the sense that they are free from constant error. Equation (2) would remain the best estimate of variability in the case of biased cues, as a constant bias preserves variability. However, the maximum gain in precision, equation (1), would not be the same (see Scarfe and Hibbard (2011) for a full treatment of biased cues). Of course, other types of bias could be present in the data, such as different constant errors for different stimulus values. A central tendency bias can cause such an effect and we discuss how to account for this later in the text.

3 It does not preserve equivalence between the optimal prediction and measured variability using both cues where the optimal prediction is calculated from the measured single cue variabilities as 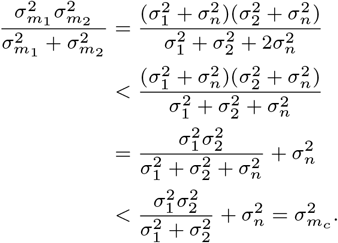 In summary, the calculated optimal prediction will imply that the observers can be more precise than is actually possible.

4 Measures of behavioural variability also lead to invalid optimal predictions. This can be seen by trying to predict the theoretical multi-cue variability from the theoretical single-cue variabilities. It can be shown that 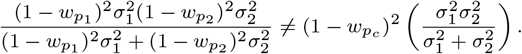

## Notes

https://osf.io/zxmpf/

### Competing Interest Statement

The authors have declared no competing interest.

### Summary of Updates

We corrected a few typos

## References

Alais, D. and Burr, D. (2004). Ventriloquist Effect Results from Near-Optimal Bimodal Integration. Current Biology, 14(3):257–262.

Ashourian, P. and Loewenstein, Y. (2011). Bayesian Inference Underlies the Contraction Bias in Delayed Comparison Tasks. PloS one, 6(5).

Aston, S., Pattie, C., Beierholm, U., and Nardini, M. (2020). Failure to account for extrinsic visual noise leads to suboptimal multisensory integration. Journal of Vision, 20(11):880.

Bejjanki, V. R., Knill, D. C., and Aslin, R. N. (2016). Learning and inference using complex generative models in a spatial localization task. Journal of Vision, 16(2016):1–13.

Berniker, M., Voss, M., and Kording, K. (2010). Learning priors for bayesian computations in the nervous system. PLoS ONE, 5(9):1–9.

Chambers, C., Sokhey, T., Gaebler-Spira, D., and Kording, K. P. (2018). The development of Bayesian integration in sensorimotor estimation. Journal of Vision, 18(12):8.

Cicchini, G. M., Arrighi, R., Cecchetti, L., Giusti, M., and Burr, D. C. (2012). Optimal Encoding of Interval Timing in Expert Percussionists. The Journal of Neuroscience, 32(3):1056 LP – 1060.

Corbin, J. C., Crawford, L. E., and Vavra, D. T. (2017). Misremembering emotion: Inductive category effects for complex emotional stimuli. Memory & Cognition, 45(5):691–698.

Crawford, L. E. (2019). Reply to Duffy and Smith’s (2018) reexamination. Psychonomic Bulletin & Review, 26(2):693–698.

Crawford, L. E., Huttenlocher, J., and Engebretson, P. H. (2000). Category Effects on Estimates of Stimuli : Perception or Reconstruction? Psychological Science, 11(4):280–284.

Duffy, S., Huttenlocher, J., Hedges, L. V., and Crawford, L. E. (2010). Category effects on stimulus estimation: shifting and skewed frequency distributions. Psychonomic Bulletin and Review, 17(2):224–230.

Duffy, S. and Smith, J. (2018). Category effects on stimulus estimation: Shifting and skewed frequency distributions—A reexamination. Psychonomic Bulletin & Review, 25(5):1740–1750.

Duffy, S. and Smith, J. (2020a). Omitted-variable bias and other matters in the defense of the category adjustment model: A comment on Crawford (2019). Journal of Behavioral and Experimental Economics, 85:101501.

Duffy, S. and Smith, J. (2020b). On the category adjustment model: another look at Huttenlocher, Hedges, and Vevea (2000). Mind & Society, 19(1):163–193.

Ernst, M. O. and Banks, M. S. (2002). Humans integrate visual and haptic information in a statistically optimal fashion. Nature, 415(6870):429–433.

Hillis, J. M., Ernst, M. O., Banks, M. S., and Landy, M. S. (2002). Combining Sensory Information: Mandatory Fusion Within, but Not Between, Senses. Science, 298(5598):1627–1630.

Hollingworth, H. L. (1910). The Central Tendency of Judgment. The Journal of Philosophy, Psychology and Scientific Methods, 7(17):461–469.

Huttenlocher, J., Hedges, L. V., and Vevea, J. L. (2000). Why do categories affect stimulus judgment? Journal of Experimental Psychology: General, 129(2):220–241.

Jamieson, D. G. (1977). Two presentation order effects. Canadian Journal of Psychology, 31(4):184–194.

Jazayeri, M. and Shadlen, M. N. (2010). Temporal context calibrates interval timing. Nature Neuroscience, 13(8):1020–1026.

Jones, S. A., Beierholm, U., Meijer, D., and Noppeney, U. (2019). Older adults sacrifice response speed to preserve multisensory integration performance. Neurobiology of Aging, 84:148–157.

Kiryakova, R. K., Aston, S., Beierholm, U. R., and Nardini, M. (2020). Bayesian transfer in a complex spatial localization task. Journal of Vision, 20(6):17.

Knill, D. C. and Saunders, J. A. (2003). Do humans optimally integrate stereo and texture information for judgments of surface slant? Vision Research, 43(24):2539–2558.

Körding, K. P. and Wolpert, D. M. (2004). Bayesian integration in sensorimotor learning. Nature, 427(6971):244–247.

Krügel, A., Rothkegel, L., and Engbert, R. (2020). No exception from Bayes’ rule: The presence and absence of the range effect for saccades explained. Journal of Vision, 20(7):15.

Laquitaine, S. and Gardner, J. L. (2017). A Switching Observer for Human Perceptual Estimation. Neuron, pages 1–13.

Murai, Y. and Yotsumoto, Y. (2018). Optimal multisensory integration leads to optimal time estimation. Scientific Reports, 8(1):13068.

Negen, J., Chere, B., Bird, L.-A., Taylor, E., Roome, H. E., Keenaghan, S., Thaler, L., and Nardini, M. (2019). Sensory cue combination in children under 10 years of age. Cognition, 193:104014.

Norton, E. H., Acerbi, L., Ma, W. J., and Landy, M. S. (2019). Human online adaptation to changes in prior probability. PLoS computational biology, 15(7):e1006681.

Odegaard, B., Beierholm, U. R., Carpenter, J., and Shams, L. (2019). Prior expectation of objects in space is dependent on the direction of gaze. Cognition, 182:220–226.

Olkkonen, M. and Allred, S. R. (2014). Short-Term Memory Affects Color Perception in Context. PlosOne, 9(1):1–11.

Olkkonen, M., McCarthy, P. F., and Allred, S. R. (2014). The central tendency bias in color perception : Effects of internal and external noise. Journal of Vision, 14(11):1–15.

Oruç, I., Maloney, L. T., and Landy, M. S. (2003). Weighted linear cue combination with possibly correlated error. Vision Research, 43(23):2451–2468.

Plummer, M. (2003). JAGS: A Program for Analysis of Bayesian Graphical Models using Gibbs Sampling. 3rd International Workshop on Distributed Statistical Computing (DSC 2003); Vienna, Austria, 124.

Rahnev, D. and Denison, R. N. (2018). Suboptimality in perceptual decision making. Behavioral and Brain Sciences, 41:e223.

Riskey, D. R., Parducci, A., and Beauchamp, G. K. (1979). Effects of context in judgments of sweetness and pleasantness. Perception & Psychophysics, 26(3):171–176.

Roberson, D., Damjanoviv, L., and Pilling, M. (2007). Categorical perception of facial expressions : Evidence for a “ category adjustment “ model. Memory and Cognition, 35(7):1814–1829.

Ryan, L. J. (2011). Temporal context affects duration reproduction reproduction. Journal of Cognitive Psychology, 23(1):157–170.

Scarfe, P. (2020). Experimentally disambiguating models of sensory cue integration. bioRxiv, page 2020.09.01.277400.

Scarfe, P. and Hibbard, P. B. (2011). Statistically optimal integration of biased sensory estimates. Journal of Vision, 11(7):12.

Sciutti, A., Burr, D., Saracco, A., Sandini, G., and Gori, M. (2014). Development of context dependency in human space perception. Experimental brain research, 232(12):3965–3976.

Tassinari, H., Hudson, T. E., and Landy, M. S. (2006). Combining Priors and Noisy Visual Cues in a Rapid Pointing Task. Journal of Neuroscience, 26(40):10154–10163.

Vilares, I., Howard, J. D., Fernandes, H. L., Gottfried, J. A., and Kording, K. P. (2012). Differential representations of prior and likelihood uncertainty in the human brain. Current Biology, 22(18):1641–1648.

